# Integrative data analysis predicts YY1 as a Cis-regulator in the 3D Cell Culture Models of MCF10A at the Stiffness Level of High Mammographic Density

**DOI:** 10.1101/365403

**Authors:** Qingsu Cheng, Mina Khoshdeli, Chongzhi Zang, Bahram Parvin

## Abstract

Previous studies have shown that in 3D cell culture models of human mammary cells (HMEC) (i) colony organizations are heterogeneous, and (ii) ERBB2 is overexpressed in MCF10A when the stiffness of the microenvironment is increased to that of high mammographic density (MD). The goal of the current study is to identify transcription factors that regulate processes associated with the increased stiffness of the microenvironment. Two HMEC premalignant lines of MCF7 and 184A1 are cultured in 3D, colonies are imaged using confocal microscopy, and colony organizations and heterogeneity are quantified as a function of the stiffness of the microenvironment. In parallel and surrogate assays, colony organizations are profiled by transcriptomics. Transcriptome data are enriched by correlative analysis with the computed morphometric indices, from 3D culture, and a subset of transcriptome data is selected. This subset is then processed with Model-based Analysis of Regulation of Gene Expression (MARGE) and publicly available ChIP-seq data to predict regulatory transcription factors. The integrative analysis indicated that YY1 regulates ERBB2 in the 3D cell culture of MCF10A when the stiffness of the microenvironment is increased to that of high MD. Subsequent experimental validation confirmed that YY1 is only expressed at the high stiffness value of the microenvironment concomitant with the overexpression of ERBB2 in MCF10A. Furthermore, using ERBB2 positive SKBR3 cell line, co-expression of YY1 and ERBB2 is absent, which indicates that YY1 regulates tumorigenicity through multiple pathways.

**Author’s summary:** MCF10A is a premalignant immortalized human mammary cell that has been isolated from a patient with fibrocystic and lost several barriers toward transformation. In an earlier study, we showed that ERBB2 is upregulated in 3D cultures of MCF10A when the stiffness of the microenvironment is increased to that of high mammographic density. Here, we leverage publicly available ChIP-seq data to predict and validate the cis-regulator of ERBB2. Our integrated experimental and computation protocol provides a pathway for elucidating regulators that can potentially be targeted for intervention.

## Introduction

We aim to identify and validate key transcription factors (TFs) and functional enhancers that regulate processes associated with the increased stiffness of the microenvironment in 3D cell culture models. Toward this objective, we integrate surrogate readouts that are generated through imaging of 3D colony formation with transcriptomic profiling data leading to a subset of the enriched gene set through correlative analysis. 3D colony formations are imaged using confocal microscopy, where each colony is segmented and represented multi-parametrically [1, 2] as a function of the increased stiffness of the microenvironment. In a parallel assay, transcriptome data are also generated, under identical conditions, and genes that correlate with computed colony (e.g., morphometric) indices are identified and enriched. Subsequently, the enriched gene set is cross-referenced with publicly available ChIP-seq data for inferring regulatory transcription factors. Although gene expression profiling, using microarray or RNA-seq techniques, can identify differentially expressed genes between two or more conditions, the gene list is not sufficient to learn about regulation. Existing methods such as ARACNE [3, 4] use transcriptomic profiling to infer cis-regulatory networks. However, these techniques require a large number of samples for expression profiling, and indirect effects, such as secondary regulatory relations, between genes, cannot be distinguished from the direct transcriptional regulatory events. ChIP-seq data measure genome-wide distributions of regulatory or epigenetic factors (transcription factors or histone modifications) that elucidate direct interactions between regulators and their target genes. By taking advantage of publicly available genomic data, including histone modification ChIP-seq and chromatin accessibility DNase-seq data from various cell types, we can predict the functional enhancer elements in the human genome.

We use a recently developed method called Model-based Analysis of Regulation of Gene Expression (MARGE) [5] to infer the cis-regulatory network. MARGE uses a compendium of published ChIP-seq data for H3K27ac (e.g., acetylation at 27th lysin residue of the Histone H3 protein) and applies a regression-based semi-supervised learning approach to predict the functional enhancers of a given gene set. The key transcription factors can then be predicted by associating the MARGE-predicted enhancer profile with thousands of existing TF ChIP-seq datasets. MARGE defines a regulatory potential for each gene by summarizing nearby H3K27ac ChIP-seq signals, and we have shown that regulatory potentials scaled across public H3K27ac profiles are more predictive of genes that are repressed or activated by BET inhibitors or the super-enhancer based approach [5]. MARGE offers three distinct advantages. First, compared with the existing master-regulator prediction methods such as super-enhancer analysis [6, 7], MARGE uses a more quantitative model to associate H3K27ac signals to each gene, and, as a result, MARGE has been shown to outperform ROSE [6, 7]. Second, MARGE uses a compendium of published ChIP-seq data for H3K27ac, a histone mark associated with the active enhancer regions in the genome [8, 9], which have been reported to be more informative for inference of gene regulation than looking at the promoters alone. Third, MARGE is designed to capture important information that can explain variations in gene expression, regardless of up/down-regulation, or what cell type or tissue origin the data come from [5]. Hence, gene regulation inference with MARGE is not limited to the availability of epigenomic data for any particular tissue type that we are studying, but can also apply to any cell system regardless of the availability of epigenomic data in the compendium.

In a recent study, we examined 3D colony organization as a function of the stiffness of the extracellular matrix (ECM) within the range of mammographic density (MD) [10]. The study concluded that ERBB2 is overexpressed in the 3D cultures of MCF10A when the stiffness of the microenvironment is increased to that of high MD in the clinical samples. The stiffness of the MD was measured on clinical histology samples using atomic force microscopy, which varied between 250 to 1800Pa [10]. Overexpression of ERBB2 was hypothesized by integrating computed morphometric indices of colony formation, from microscopy readouts, with the matched gene expression data and then validated using immunofluorescence staining and microscopy. Although we have shown that the relationship between ERBB2 and stiffness of the microenvironment is specific to the 3D cell culture model and MCF10A, the regulators of ERBB2 are the primary motivation of this study. The details of the experimental design, microscopy, and computational analysis are included in the material and method section, i.e., supplementary files for online publication.

## Results

### The frequency of aberrant phenotypes are increased as a function of increased stiffness

Two human mammary epithelial cell (HMEC) lines were cultured in 3D, and the stiffness of the microenvironment was modulated between 250 to 1800 Pa., which is the stiffness values at low and high mammographic density [10]. At each stiffness value, colony formation was imaged using confocal microscopy, and from the same passage, RNA was collected for transcriptomic profiling. Colony formation indices were quantified using BioSig3D [1, 2], where for each colony a number of indices are computed. These indices include geometrical and morphological features of colony formation such as flatness, elongation, colony size, lumen formation, and changes in the proximity of nearby cells. The rationale being that aberrant colony organization (e.g., colony flatness) is one of the first indicators of dysplasia and an early marker of malignant progression. One of the capabilities of the BioSig3D is that heterogeneity can be profiled through consensus clustering, where the frequency of occurrence of each cluster can then be quantified. As a result, sensitive assays can be developed and profiled. One example of heterogeneity analysis is shown in **Fig. 1**, where elongation is shown to have two subpopulations, and the frequency of occurrence of the aberrant subpopulation increases as a function of the stiffness of the microenvironment. The frequency of aberrant colony formation is consistent for both cell lines. Another example is shown in **Fig. 2**.

**Figure 1.**
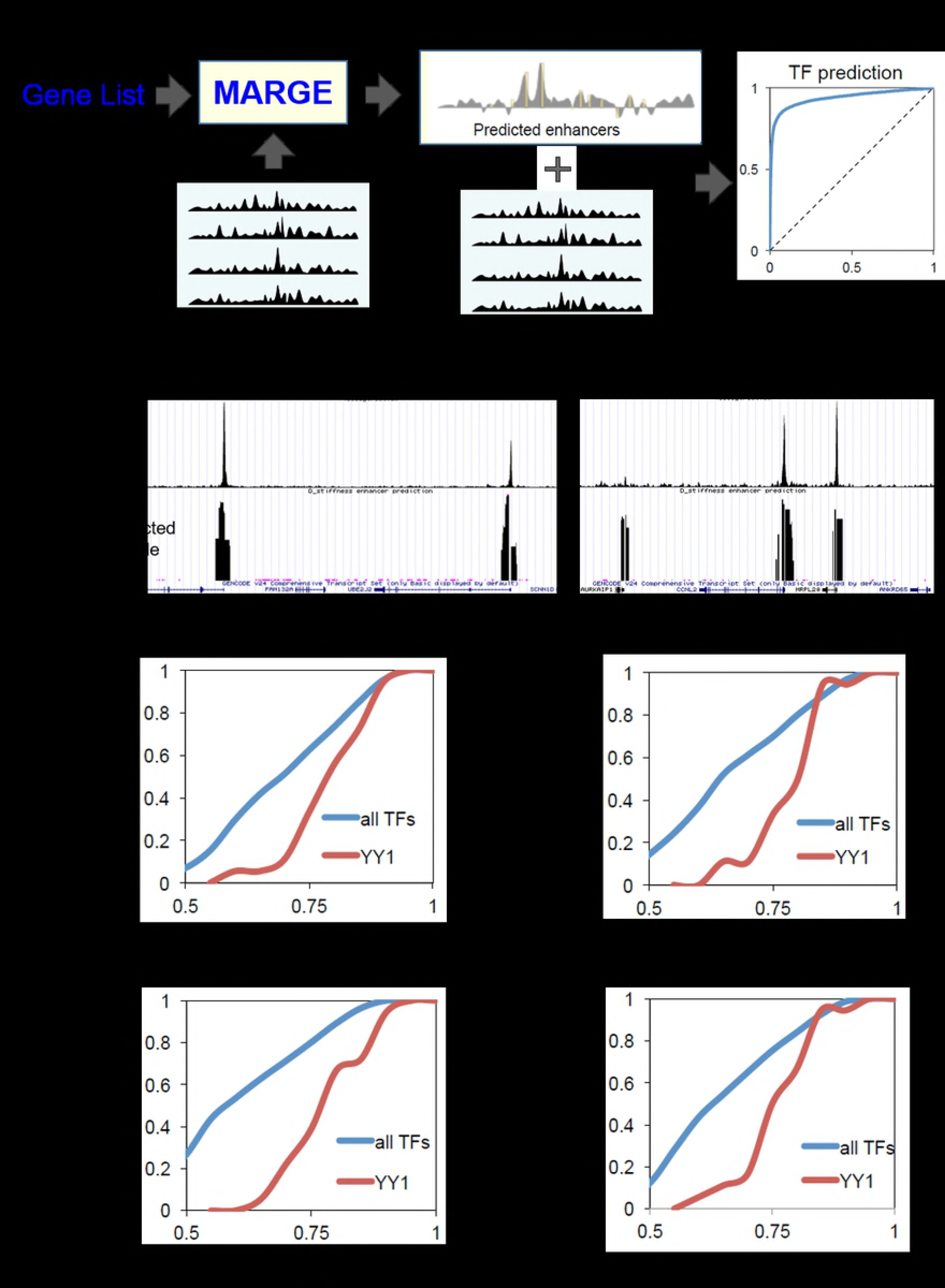
Colony shapes are heterogeneous: (a) Consensus clustering identifies two morphometric subtypes quantified with the colony elongation index as a function of increased stiffness of the microenvironment from 250 to 1800Pa. (b-c) Representative of colony organization for each of the subtypes imaged through confocal microscopy with each nucleus segmented with BioSig3D. (d) Frequency of occurrence of colony elongation increases as a function of increased stiffness of the microenvironment. Scale bar is 20 µm.

### Proliferation rate, within each colony, is cell line dependent

Phenotypic analysis of the colony size as a function of increased stiffness inferred two subtypes for each of the cell lines as shown in **Fig. 2(a-b)**, where the colony size is profiled by the number of cells in each colony. The results indicate that for MCF10A, there is a dominant population of smaller colony size at the low stiffness of the microenvironment, but the frequency of larger colony size increases as a function of the higher stiffness of the microenvironment. On the other hand, for 184A1 (http://hmec.lbl.gov/mindex.html from Stampfer Lab), two equal subpopulations are present at the low stiffness of the microenvironment, but the frequency of larger colony size decreases as a function of the increased stiffness of the microenvironment. These data suggest that (i) MCF10A is more clonal and homogenous than 184A, (ii) at the high stiffness of the microenvironment, both cell lines behave similarly, and (iii) subtypes of colony size can be used to fingerprint each cell line.

**Figure 2.**
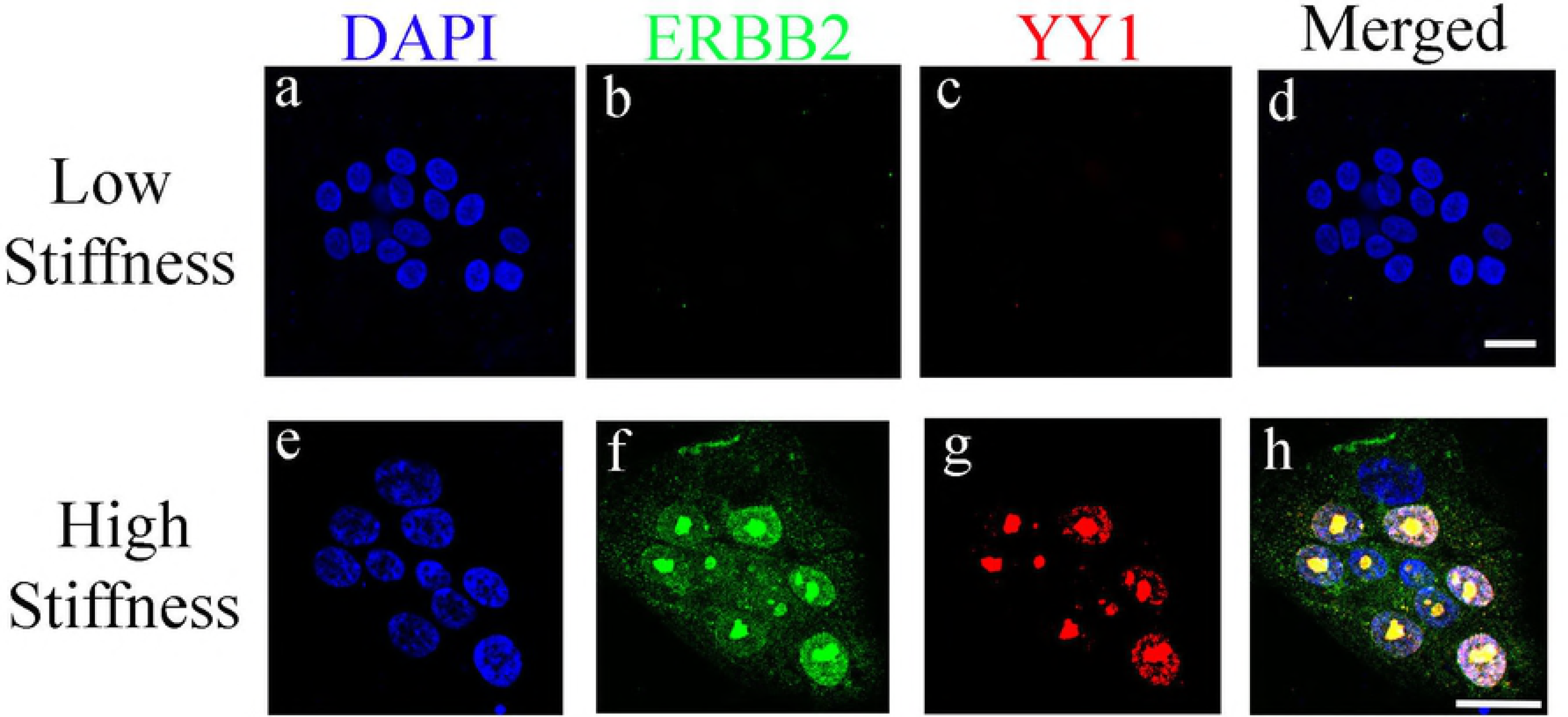
Colony proliferation rates are heterogeneous. (a) Consensus clustering identifies two diverse morphometric subtypes for MCF10A with the representative of each subpopulation shown. The frequency of larger colonies increases as a function of increased stiffness of the microenvironment. (b) Consensus clustering identifies two approximately equal morphometric subtypes for 184A1 with the representative of each subpopulation shown. The frequencies of smaller colonies increases as a function of increased stiffness of the microenvironment. Scale bar is 20 µm.

### YY1 is predicted as a regulatory transcription factor for MCF10A and 184A1

For each of the cell lines, genes that correlate with phenotypic indices (e.g., colony elongation, colony size) were identified and a subset of genes was selected based on their p-values. The correlative analysis enriches the gene expression data with the computed colony formation indices, which serve as a surrogate assay. The raw data and the enriched gene sets are included as data file for on-line publication. The enriched gene sets were computed for two colony formation indices, applied to MARGE [5] and the relevant publicly available H3K27ac ChIP-seq datasets, and the functional enhancer-profiles that regulate these genes are predicted, as shown in **Fig. 3a.** The relevant H3K27ac ChIP-seq datasets, selected by MARGE, include samples from the breast cancer cell line MCF-7, which indicates biological similarity in gene regulation profiles between the cell lines, as shown in **Supplementary Table 1**. An example of the MARGE-predicted enhancer profile is shown in **Fig. 3b.**

We then investigated the association between the MARGE-predicted functional enhancer profile and 2264 ChIP-seq datasets collected from the public domain [13] representing genomic profiles of over 400 TFs in various cell types. Among these TFs, with the available ChIP-seq data, Yin-Yang 1 (YY1) was identified as highly associated with the MARGE-predicted enhancer profile in both cell lines. The analysis is based on the criteria that YY1 binding sites in most YY1 ChIP-seq datasets align with the top-ranked enhancers in the genome with a P-value less than 0.001, as shown in **Fig. 3c-f**. It is important to note that YY1 was predicted with only *15* human breast samples of H3K27ac ChIP-seq. YY1 is a ubiquitous transcription factor that plays an important role in normal biological processes such as differentiation and proliferation, as well as tumorigenesis [15], where tumorigenesis is induced through cell cycle regulation, inhibition of the p53 tumor suppressor gene, or regulation of apoptosis. YY1 has been shown to cooperate with the AP-2 transcription factor (cell growth and differentiation) to stimulate ERBB2 [16], which is generally overexpressed in breast cancer and is negatively regulates p27 (a cell cycle inhibitor) [17]. p27 is known to function as a tumor suppressor, which is rarely mutated in human cancers. Increased stiff of the matrix leads to Rho activation [18], where Rho activation has been linked to increased proliferation [19] and is necessary to remove the inhibition of cell cycle progression that is imposed by p27. Furthermore, inhibition of YY1 in MCF7 has been shown to transform 3D colony formation to that of MCF10A [17]; however, 3D colony formation has been cell line dependent with the down-regulation of YY1, which is potentially due to the degree of malignant properties for each of the cell line.

**Figure 3.**
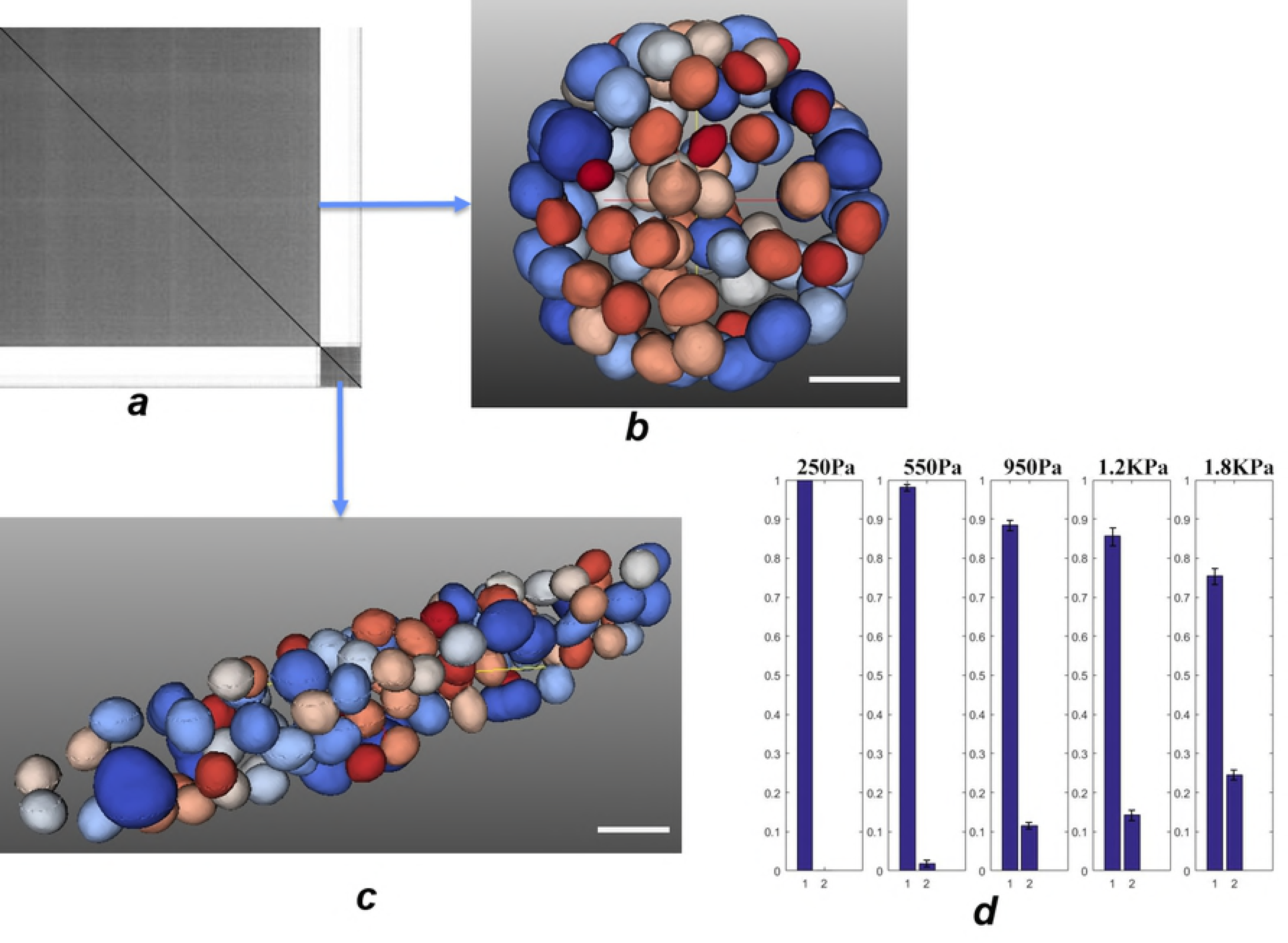
MARGE predicts YY1 as a functional transcription factor (TF) regulating phenotype-associated genes. (a) MARGE schematic. MARGE predict functional cis-regulatory (enhancer) profiles of a given gene list, using a compendium of public H3K27ac ChIP-seq profiles. Key TFs are further predicted by associating the cis-regulatory (enhancer) profiles generated by MARGE with each of 2264 collected public TF ChIP-seq datasets. For each TF dataset, a receiver operating characteristic (ROC) curve is generated, and the area under the curve (AUC) is used to assess the association of the regulatory profile with this TF in the genome. (b) Sample genome browser snapshot of a public YY1 ChIP-seq profile (upper track) and an MARGE-predicted cis-regulatory (enhancer) profile (lower track). YY1 ChIP-seq dataset is from K562 cell line (Pope, et al., 2014). The enhancer profile is predicted from genes correlated with stiffness. (c-f) YY1 is highly enrich regardless of the phenotypic indices for MCF10A. Cumulative distributions of AUC scores for predicting all 2264 ChIP-seq datasets for different TFs’ binding (blue) and 18 datasets for YY1 binding (red). Having significantly higher AUC scores than the all-TF background, YY1 is predicted as a functional regulator of the input gene sets for cell line MCF10A. (P-value < 0.001, by K-S test.)

### YY1 activates ERBB2 in MCF10A at high stiffness values of the microenvironment

In a recent paper, we showed that ERBB2 is overexpressed in 3D cell culture models of an MCF10A cell line by simply increasing the stiffness of the microenvironment within the range of MD, i.e., from 250 to 1800Pa [10]. Having identified YY1 as a TF regulator using MARGE, we then designed three experiments with the necessary set of controls for validating overexpression of YY1 in 3D cell culture models. To begin, control cells were engineered by overexpressing and knocking down YY1 using the CRISPR technology with a sample result shown in **Supplementary Fig. 1** and **2**. The first experiment was performed with MCF10A, which indicated that (i) YY1 regulates ERBB2 at high stiffness, (ii) if YY1 is turned off with CRISPR, then ERBB2 is never activated either at low or high stiffness values of the microenvironment, and (iii) if YY1 is upregulated then ERBB2 is also activated even at low stiffness of the microenvironment. **Fig. 4** shows co-localization of YY1 with ERBB2. The second experiment was performed with the non-transformed 184A1 HMEC; however, neither overexpression of ERBB2 nor YY1 were observed. The third experiment was performed with the ERBB2 positive cell line of SKBR3, where control cells were engineered to overexpress and knock down YY1. Regardless of the stiffness of the microenvironment and YY1 being on up or down, ERBB2 was always expressed in this cell line.

**Figure 4.**
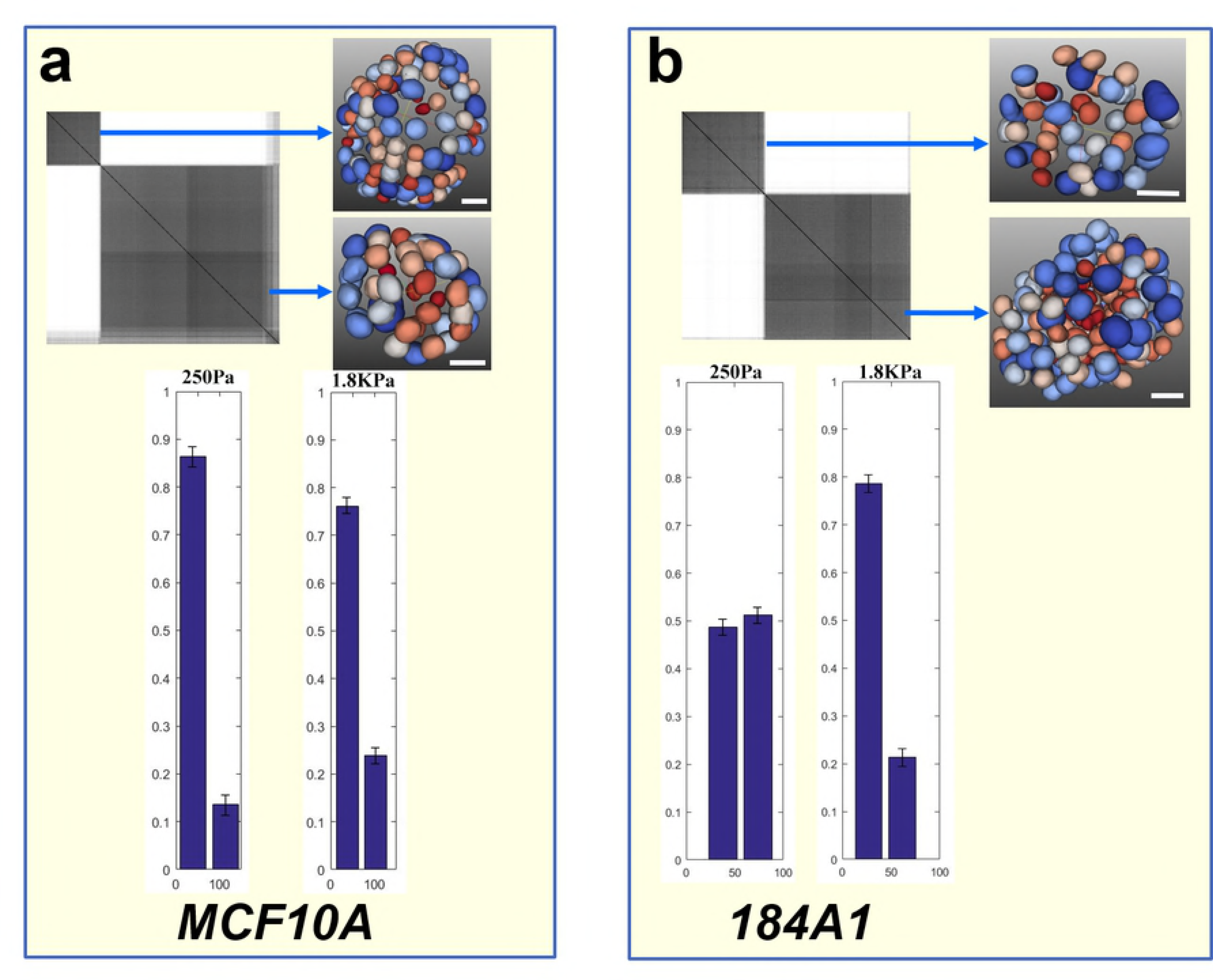
YY1 regulates ERBB2 in MCF10A monitored by fluorescent microscopy. A cross section of a multicellular system imaged with confocal microscopy. 3D colony formation is fixed at day 10 and stained. Top row shows a colony with DAPI, ERBB2, YY1 stain at 100 Pa. Bottom row shows DAPI, ERBB2, YY1, and merged signal of YY1 and ERBB2 at 1800Pa. Scale bar is Scale bar is 20 µm.

## Discussion

The following observations are made using the two human mammary epithelial cell lines of MCF10A and 184A1. These two cell lines were selected because, at high stiffness, MCF10A expresses ERBB2 while 184A1 does not [10]. Furthermore, MCF10A originates from a patient with the fibrocystic *disease*, while 184A1 originates from the normal patient following reduction mammoplasty. We conclude that (I) Increased stiffness of the microenvironment, within the range normal mammographic density, also increased the frequency of aberrant colony formation for both cell lines. (II) Proliferation indices can be used to assess diversity and heterogeneity in each of the cell lines, and subtyping based on computed indices has the potential to identify each cell line uniquely and stratify cell lines phenotypically. (III) Having enriched the gene expression data with computed colony indices, MARGE predicted YY1 to be highly enriched as one of the cis-regulatory hubs for both cell lines. It is important to note that neither ERBB2 nor YY1 are enriched at the transcription level. (IV) YY1 is co-expressed with ERBB2 in MCF10A when the stiffness of the microenvironment is increased to that of high mammographic density, but the co-expression of ERBB2 and YY1 in 184A1 is absent. However, in an ERBB2 positive cell line of SKBR3, overexpression or knock out of YY1 had no impact on the expression of ERBB2. Additional observations suggest overexpression of YY1 in SKBR3 and a number of other tumor cell lines [17]. Similar observations were also made when the culture was performed in the presence of lactacystin, a protease inhibitor, and a nonsteroidal anti-inflammatory drug (NSAID) [20]. However, both of these studies have used the 2D cell culture models of the cell lines and bulk measurement. It is well known that the patterns of gene and protein expression are differentially expressed between the 2D and 3D cell culture models. For example, our analyses have shown that (a) ERBB2 is only expressed in 3D cell culture models at the stiffness value of high mammographic density [10], and (b) PPARγ, one of the biomarkers of invasive phenotypes is only expressed in 3D and not in 2D [21].

In conclusion, integrative morphometric and transcriptomic data has enriched the molecular signature for inference of the cis-regulatory networks. Our integrative analysis suggests that YY1 upregulates ERBB2 in the 3D cell culture of MCF10A when the stiffness of the microenvironment is increased to that of high mammographic density. In addition, YY1 is also highly enriched in 184A1, but not expressed within the same range of the experimental conditions. This is possible because MCF10A is more differentiated and is derived from a benign proliferative breast tissue, which is spontaneously immortalized without defined factors [22]. Furthermore, using ERBB2 positive SKBR3 cell line, co-expression of YY1 and ERBB2 is absent, which indicates that YY1 regulates tumorigenicity through multiple signaling pathways. Similar analyses are applicable for the analysis of a wide array of biological processes and cell signaling.

## Methods

### Embedded 3D culture model

Embedded method was used for the 3D cell culture model[11]. For each stiffness condition as per mammographic density (MD)[12], three replicates of 3D cell cultures of MCF10A human mammary epithelial cells (HMEC) were performed. The stiffness of the microenvironment per MD was modulated by varying concentration of agarose, 1% agarose for low MD and 2.0% agarose for High MD. The initial seeding density for 3D cell culture is 8000 cells per well. The procedure of preparing 3D culture was described as following. The cells were carefully suspended in μL Matrigel matrices (BD). Next, 40 μL of 2.0% and 4% agarose solution (Sigma) were added and thoroughly mixed. The formed gels have final concentrations of agarose at 1.0% and 2.0%. The corresponding stiffnesses were 250 and 1800 Pa, respectively. The mixtures were placed in an 8-well chambered coverglass (Nunc Lab Tek II), and incubated for 20 min, allowing each matrix to gel before adding the growth medium (500 μL/well). Cells were maintained in culture for 8 days while changing the medium once every third day. Changing of the medium was also monitored with phase microscopy for proper growth and colony formation.

### Cell subculture and maintenance

MCF10A and SKBR3 cultured sequentially following an initial pilot project, validation, and hypothesis generation. Cells were maintained in a 37°C incubator with 5% CO_2_. The MCF10A model system was obtained from ATCC at passage > 50 and was then subcultured (P5-P7) for the proposed studies. MCF10A were cultured in DMEM/F12 medium (Life Science Technology) supplemented with 5% horse serum (Life Science Technology), 10 ng/mL EGF (Life Science Technology), 50 ng/mL chlora toxin (Sigma), 200 μg/mL insulin (Sigma), 100 ng/mL hydroxylcortisone (Sigma), and 1% penicillin-streptomycin (Life Science Technology). SKBR3 were cultured in DMEM supplemented with 10% of FBS (Life Science Technology) and 1% penicillin-streptomycin.

### Transfection

The CRISPR-cas9 technology was used to transfect MCF10A cell lines to overexpress/knockout YY1 (Santa Cruz Biotechnology, SC400295/SC409112). The vendor’s protocol was followed without any modifications as follows: (I) 10,000 cells were seeded per well in a 12-well plate tissue culture plate and incubated in 2 mL of antibiotic-free standard growth medium. (II) Cells were monitored for their confluency to conclude healthy and proliferating cells, which are required for CRISPR Activation Plasmid transfection, i.e., confluency should be approximately 60-80. (III) 300 µl YY1 activation Plasmid Transfection Reagent Complex (Complex component details were described in company provided protocol was added dropwise to each well. (IV) The plate was gently mixed by swirling, and cells were incubated for hours under conditions normally used to culture the cells. (V) Empty plasmid vectors were set aside as a negative control. (VI) After 24 hours, transfected cells were stained with the YY1 antibody for validation and subsequently transferred to the 3D culture for further testing.

### Fixation and immunofluorescence staining

Cell cultures in the hydrogel and on the plastic plate were fixed at room temperature in 10% formalin for 30 min. After 3 times of PBS washes, cells were permeabilized using a 0.1% Triton X-100 solution for 5 min. Cell cultures were washed 3 times with PBS, blocked using a 1% bovine serum albumin (BSA) solution (Sigma) supplemented with 1% goat IgG (Jackson ImmunoResearch), and the blocking solution was maintained for 1 h. In some cases, samples were stained for a specific molecular endpoint, but in all cases, DAPI nuclear stain was applied for profiling colony organization. Monoclonal rabbit anti-human primary antibodies (*e.g*., ERBB2 EP1045Y, and YY1 EPR4652) (Abcam) were diluted in blocking solution (1:500), incubated for 1 h, and followed by 3 PBS washes. Goat anti-rabbit secondary antibody conjugated with Alexa and Alexa 565 (A11008 and A11011) (Life Science Technology) was diluted in the blocking solution (1:250), conjugated to primary antibody for 1 h, and followed by 3 PBS washes. Subsequently, nuclei were stained with 4’-6-diamidino-2-phenylindole (50 ng/mL) (DAPI D1306, Life Science Technology).

### Fluorescent microscopy

Stained samples were imaged using a Zeiss LSM 710 system equipped with a Zeiss Apochromat 40X/1.1 (0.8 mm WD) water immersion objective lens. The excitation filters were set at 405, 488 and 568 nm, and the emission filters were set to receive signals between 420-480, 420-550 and 597-700 nm, respectively. The laser intensity was set at 1%. A twin-gate main beam splitter with two wheels, each wheel having 10 filter positions (and thus 100 possible combinations), was used to separate excitation and emission beams. The pinhole was set at “1”. The digital gain was adjusted to approximately ¾ of the maximum gain, which kept the dynamic range of the pixels between 500-2000 (12 bits). The 3D stack function of the Zen software was used to collect raw 3D information for each colony and was concurrent with “Z correction” of the pixel intensity for thick samples. The voxel size was set to 0.25µm×0.25µm×1µm, the image files were saved as lsm files, and uploaded to the BioSig3D imaging bioinformatics system [1].

### RNA analysis

Total RNA collected from different stiffness, 250 Pa to 1800 Pa, were extracted using a Qiagen RNAeasy micro kit, following the in-kit protocol. The RNA quality was checked to assure that RNA integrity number (RIN) was over 7, out of maximum 10. Total RNA extractions were sent to the UCLA UNGC core facility for profiling, where UCLA performed their own quality control check independently. Subsequently, UCLA amplified the total RNA by Ambion amplification and profiled gene expression on Illumina human ht-12 chip in triplicates. The chips were scanned, gene expression data were normalized, and p-values for the quality of each gene was computed, and a spreadsheet was delivered for integrative analysis.

### Enhancer and key TF prediction using MARGE

MARGE (version 1.0) was used to predict the genomic enhancer profile using the highly correlated gene set as input. To obtain the best enhancer prediction, MARGE regression was executed under default parameters for 10 times and the enhancer profile with highest prediction evaluation measure (AUC) was selected. The predicted enhancer profile is a ranked list of ~2.7 million cis-elements (candidate enhancers) in the human genome, obtained as a union set of all DNase hypersensitive sites from over 400 human DNase-seq datasets. 2264 ChIP-seq datasets were collected from the human genome for TFs and chromatin regulators that is also referred to as “factor”[13]. For each factor, MACS2[14] was used to identify the binding sites under a false discovery rate (FDR) cutoff of 0.01 and five fold-enrichment cutoff. A present/absent assignment was placed on each of the 2.7 candidate enhancer sites to show whether a site is bound by this factor or not that is determined by peak region overlap. Subsequently, a receiver operational curve (ROC) is generated using the ranked enhancer prediction of each factor, and the area under the ROC curve (AUC) is calculated to evaluate the association of this factor. Finally, the AUC of 18 YY1 ChIP-seq datasets were compared with the null distribution of AUC of all 2264 ChIP-seq datasets and a p-value was calculated under the K-S test.

## List of Abbreviations

3D: three dimensional
ECM: extracellular matrix
YY1: yinyang
HMEC: human mammary epithelial cell
ERBB2: Receptor tyrosine-protein kinase erbB-2
TF: transcriptional factor
MD: mammographic density
MARGE: model based analysis of regulation of gene expression
PPARγ: peroxisome proliferator activated receptor γ
EGF: epidermal growth factor
CRISPR: clustered regularly interspaced short palindromic repeats

## Declaration

No human or animal tissues were used in this study.

## Declaration subheading

NA

## Consent to publish

All authors have read the manuscript and forward their consent to publication.

## Competing interest

The authors declare no financial competition.

## Funding

This work has been supported by NIH R01CA140663 (BP) and K22CA204439 (CZ).

## Availability of data and materials

All gene expression data have been uploaded as a part of the Supplementary Materials. These include raw data and correlative analysis.

## Author contributions

BP and QC designed the study. QC performed cell culture, immunofluorescent staining and image acquisition. MK performed correlative studies of gene expression with the morphometric data and morphometric subtyping. CZ performed MARGE study. BP, QC, and CZ wrote the manuscript. All authors read and approve the manuscript.

## Supporting information Legends

Supporting Information. SI1: Supplementary Table and Figures

Supporting Information. SI2: Raw gene expression data as a function of stiffness of the microenvironment

Supporting Information. SI3: List of genes sorted by their p-value against computed flatness in MCF10A.

Supporting Information. SI4: Keys for Gene expression data

